# FAP-retargeted Ad5 enables *in vivo* gene delivery to stromal cells in the tumor microenvironment

**DOI:** 10.1101/2022.12.19.520931

**Authors:** K. Patricia Hartmann, Merel van Gogh, Patrick C. Freitag, Florian Kast, Gabriela Nagy-Davidescu, Lubor Borsig, Andreas Plückthun

## Abstract

Fibroblast activation protein (FAP) is a cell surface serine protease that is highly expressed on reactive stromal fibroblasts, such as cancer-associated fibroblasts (CAFs), and generally absent in healthy adult tissues. FAP expression in the tumor stroma has been detected in more than 90% of all carcinomas, rendering CAFs excellent target cells for a tumor site-specific adenoviral delivery of cancer therapeutics. Here, we present a tropism-modified human adenovirus 5 (Ad5) vector that targets FAP through trivalent, designed ankyrin repeat protein (DARPin)-based retargeting adapters. We describe the development and validation of these adapters via cell-based screening assays and demonstrate adapter-mediated Ad5 retargeting to FAP^+^ fibroblasts *in vitro* and *in vivo*. We further show efficient *in vivo* delivery and *in-situ* production of a therapeutic payload by CAFs in the tumor microenvironment (TME), resulting in attenuated tumor growth. We thus propose using our FAP-Ad5 vector to convert CAFs into a ‘biofactory’, secreting encoded cancer therapeutics into the TME to enable a safe and effective cancer treatment.

## Introduction

Over the last few decades, cancer therapy has benefited substantially from immunotherapeutic approachess involving monoclonal antibodies (mAbs) and recombinant cytokines with anti-tumor activity. However, systemic application of such agents is often still restricted due to severe dose-limiting toxicities as well as off-target effects, both hampering therapeutic efficacy and safety^1, 2^. One promising strategy to overcome these limitations is to deliver the genetic information for the *in-situ* production of the effector proteins utilizing suitable delivery systems such as viral vectors. One of the most commonly used viral vector is derived from human adenovirus serotype 5 (Ad5), a non-enveloped double-stranded DNA virus consisting of three major capsid proteins: the hexon, penton, and fiber^3, 4^. In comparison to other viral vectors, mainly derived from lentivirus (LV) or adeno-associated virus (AAV), Ad5-based vectors bring the following superior characteristics for clinical use: (i) they do not integrate into the host cell genome and are therefore safe, (ii) they efficiently transduce dividing and non-dividing cells, and (iii) they have a large packaging capacity of up to 37 kilobase pairs (kb)^5^. The large packaging capacity allows for a delivery of multiple payload genes and renders adenoviral vectors very attractive for combination cancer immunotherapy.

Despite all favorable features, tissue-specific delivery of Ad5-based vectors is still compromised *in vivo*. One hurdle is the strong liver tropism of Ad5, especially upon systemic administration, mainly resulting from interactions of the hexon’s hypervariable regions (HVRs) with host molecules, causing vector sequestration by liver-resident cells^6-8^. Another limitation of Ad5 vectors is represented by the virion’s natural cell specificity and difficulties in efficiently redirecting the vector to the desired tissue. To overcome these challenges, binding to cellular receptors that are naturally involved in viral cell entry needs to be prevented, while introducing specificity for cell surface molecules located on the cell type of choice. Cell entry of Ad5 occurs in sequential steps: first, the knob domain of the fiber protein binds to its high-affinity, primary attachment receptor termed coxsackievirus and adenovirus receptor (CAR)^9, 10^. Then, an arginine-glycine-aspartate (RGD) motif in the penton base binds to integrins on the cell surface, which initiates the cellular virus uptake by clathrin-mediated endocytosis^11, 12^.

Previous retargeting efforts have aimed at modifying Ad5 interactions through various manipulations of the viral capsid, including genetic mutations, chemical modifications, and genetic fusions of targeting peptides mainly to the fiber protein^13-17^. However, these attempts have been accompanied with reduced transduction efficiency and insufficient vector targeting. To address these drawbacks, we have previously developed a generic, protein-based de- and retargeting platform for Ad5 vectors consisting of two parts: (i) a hexon-binding, single-chain variable fragment (scFv)-based trimeric shield that protects the vector from undesired host factor interactions accounting for its *in vivo* liver tropism^18^. Additionally, this shield protects the vector from immune clearance, thereby helping to overcome yet another hurdle, namely the widespread pre-existing humoral immunity against Ad5 in the human population^19, 20^. (ii) A bispecific, designed ankyrin repeat protein (DARPin)-based modular adapter binding trivalently to the fiber knob, thereby preventing CAR-interactions and thus reducing CAR-mediated cell transduction, while allowing to introduce a defined cell specificity by incorporating appropriate targeting moieties^21-23^. Importantly, this bispecific adapter does not require genetic fusion to the fiber protein and can hence be applied to any Ad5-derived vector, including high-capacity Ad5 vectors.

By applying DARPin adapters specific for tumor cell markers, such as human epidermal growth factor receptor 2 (HER2), to a shielded HVR7-mutated Ad5 vector, we recently demonstrated efficient vector detargeting from the liver and other organs, as well as successful vector retargeting to tumor cells *in vivo*^18^. Following the validation of our <u>sh</u>ielded and <u>re</u>targeted <u>ad</u>enoviral (‘SHREAD’) delivery platform, we aim to expand our retargeting system to stromal cells in the tumor microenvironment (TME) to enable alternative vector targeting and tumor treatment strategies. One of the most abundant stromal cell types in the TME are cancer-associated fibroblasts (CAFs), which evolve alongside cancer cells during tumor development and progression^24, 25^. Importantly, CAFs express the cell surface glycoprotein fibroblast activation protein α (FAP) that is selectively present in reactive stromal fibroblasts and generally absent in healthy adult tissues^26, 27^. More importantly, FAP on CAFs was found to be overexpressed in more than 90% of all carcinomas, including lung, colorectal, prostate, and breast cancer^28^, and was therefore established as a promising target for tumor therapy, imaging, and diagnosis^29-42^. These characteristics render CAFs an attractive cell type for adenoviral vector retargeting via FAP, to then utilize CAFs as a ‘biofactory’ for the local production and secretion of cancer therapeutics in the TME.

Here, we present a cell-based screening approach following ribosome display selections to generate FAP-specific DARPin adapters that enable retargeting of Ad5-derived vectors to fibroblasts via FAP. We describe the characterization of the retargeting adapters, and then provide *in vitro* and *in vivo* data demonstrating adapter-mediated Ad5 retargeting to FAP^+^ fibroblasts. Furthermore, we show efficient *in vivo* delivery of a therapeutic payload to FAP^+^ fibroblasts in the TME resulting in reduced tumor growth. These data prove feasibility of our concept: to deliver genes encoding biomolecules with anti-tumor activity through a FAP-retargeted Ad5 to CAFs in the tumor stroma.

## Results

### Ribosome display selection of DARPins against hFAP

FAP is a 170 kilodalton (kDa) type-II transmembrane serine protease with a large C-terminal extracellular domain containing the enzyme’s active site^43^. To redirect our Ad5 vector to CAFs via FAP, we built on our previously developed modular adapter consisting of (i) a retargeting DARPin mediating cell-specificity, (ii) the fiber knob-binding DARPin 1D3, and (iii) the trimerizing protein SHP derived from lambdoid phage 21 (Fig. 1a). To generate the retargeting DARPin with specificity for FAP, we used recombinant human FAP (hFAP) and performed ribosome display selections as described before^44^. A total of four selection rounds yielded 380 DARPin candidates, which were subsequently analyzed for hFAP binding. Notably, these candidates contained mainly two (N2C) or three (N3C) internal repeats with randomized surface flanked by capping repeats.

**Figure 1:**
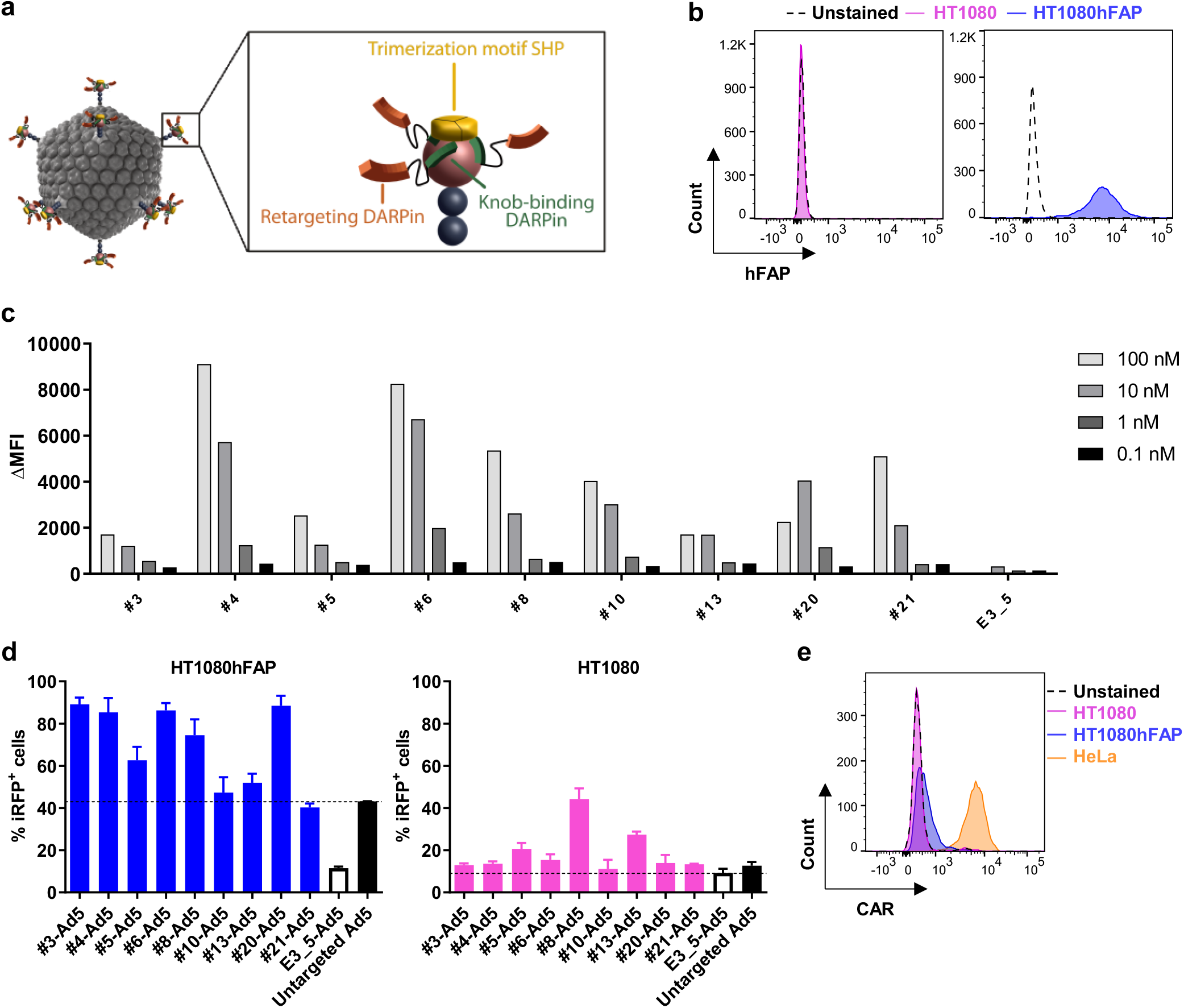
Generation and validation of DARPin-based hFAP-specific adenoviral retargeting adapters. **(a)** Schematic representation of a bispecific trimeric DARPin adapter for adenoviral retargeting. The retargeting DARPin (orange) with specificity for a selected cell surface molecule (e.g., FAP) is fused via a long flexible linker to the knob-binding DARPin 1D3 (green) that in turn is fused via a short linker to the trimerizing protein SHP from lambdoid phage (yellow). The bispecific trimeric DARPin adapter forms a highly stable clamp around the fiber knob (red) to block natural cellular interactions and redirect adenoviral tropism to selected cells (e.g., FAP^+^ cells). **(b)** Flow cytometry analysis of hFAP expression of the parental HT1080 and HT1080hFAP cell line upon hFAP antibody staining. **(c)** Cell-based adapter binding assay on target and non-target cells. The purified ‘Top 9’ adapters constructed with the selected hFAP-specific DARPins were titrated on hFAP^+^ HT1080hFAP and hFAP^−^ HT1080 cells in the concentration range of 0.1 – 100 nM. Binding was detected via flow cytometry by staining of the His-tagged adapter, and specific binding signals were determined as ΔMFI = MFI (HT1080hFAP cells) - MFI (HT1080 cells). The non-binding control adapter E3_5 was applied as a negative binding control. Bars represent specific binding signals of single-point measurements. Representative data of two independent experiments are shown. MFI = Mean fluorescent intensity. **(d)** Transduction of target and non-target cells by hFAP adapter-retargeted Ad5. Recombinant Ad5 encoding iRFP670 was pre-incubated with the ‘Top 9’ hFAP adapters (colored filled bars) or the E3_5 blocking adapter (black empty bar) and tested for transduction of hFAP^+^ HT1080hFAP and hFAP^−^ HT1080 cells in comparison to the untargeted Ad5 (black filled bar) at a multiplicity of infection (MOI) of 20 (plaque-forming units (PFU)/cell). Transduction was assessed from cellular expression of iRFP670 detected by flow cytometry. Dashed lines indicate cut-off levels above which functional and hFAP-specific adapters were identified. Bars represent mean transduction level of two biological replicates ± standard deviation (SD). Representative data of three independent experiments are shown. **(e)** Flow cytometry analysis for CAR expression of the HT1080 and HT1080hFAP cell line in comparison to the positive control HeLa cell line upon CAR antibody staining.

To allow a high-throughput binding analysis, we established a flow cytometry-based cell-binding assay using suitable target and non-target cell lines. The parental HT1080 fibrosarcoma cell line (presenting no endogenous hFAP) and a stably transfected clone with high expression of hFAP, termed ‘HT1080hFAP cell line’^39^, were used (Fig. 1b). We screened all 380 DARPins for hFAP binding on cells, using crude *Escherichia coli* (*E. coli*) extracts, and identified 91 putative binders, testing first the target cell line only (Supplementary Fig. 1a). Then, this subpopulation was assessed for binding on both the target and non-target cell line, revealing 25 specific binders as determined by the ratio of specific and unspecific binding signal, the latter set by the unselected, non-binding DARPin E3_5^45^ (Supplementary Fig. 1b). Sequencing of the 25 specific binders identified 20 unique binders with high diversity, of which eleven were N3C and nine were N2C proteins.

We expressed the selected 20 binders in small-scale *E*.*coli* cultures and purified them using immobilized metal affinity chromatography (IMAC). We confirmed the correct molecular mass via sodium dodecyl sulfate polyacrylamide gel electrophoresis (SDS-PAGE) (Supplementary Fig. 1c), and applied analytical size exclusion chromatography (SEC) to select monodisperse proteins (Supplementary Fig. 1d). Additionally, we re-tested the purified proteins for hFAP binding on target and non-target cells, leading to the selection of ten well-behaved, hFAP-specific DARPins (Supplementary Fig. 1e). Amongst these ten DARPins, particularly candidate #6 yielded very high binding signals even at subnanomolar concentrations, thereby outperforming all other selected DARPins. To further investigate binding to hFAP, we measured the binding kinetics by surface plasmon resonance (SPR) for this DARPin and determined a K_D_ value of 163 pM (Supplementary Fig. 2).

**Figure 2:**
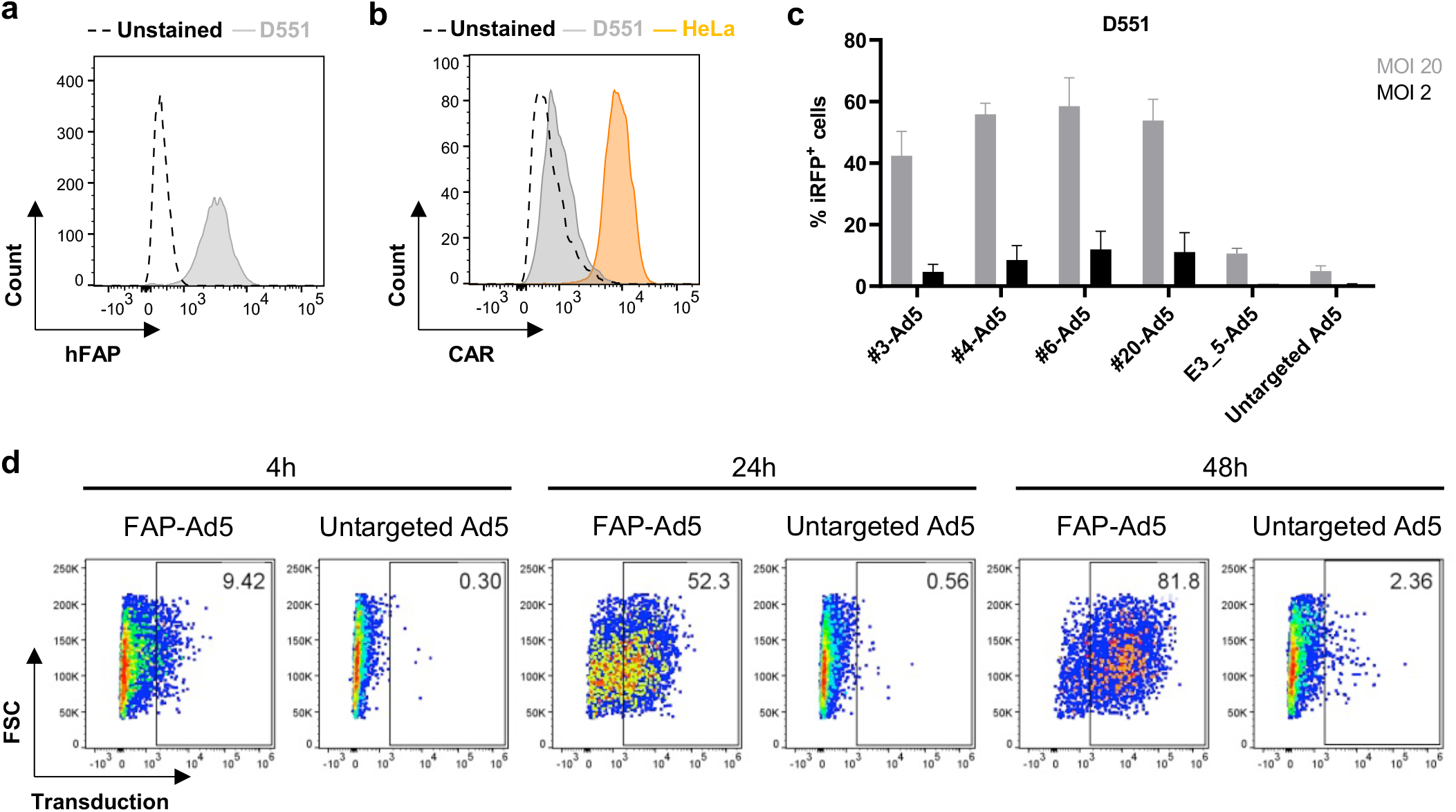
Selected hFAP adapters mediate adenoviral transduction of hFAP-expressing human fibroblasts. Flow cytometry analysis for **(a)** hFAP expression and **(b)** CAR expression of the human fibroblast D551 cell line upon antibody staining. The HeLa cell line served as a positive control in the CAR expression analysis. **(c)** Transduction of human fibroblasts by hFAP adapter-retargeted Ad5. Recombinant Ad5 encoding iRFP670 was pre-incubated with the selected ‘Top 4’ hFAP adapters or the E3_5 blocking adapter and analyzed at two different MOIs (PFU/cell) for transduction of hFAP^+^ D551 cells in comparison to the untargeted Ad5. Transduction levels were determined via cellular expression of iRFP670 detected by flow cytometry. Bars represent mean transduction level of two biological replicates ± SD. Representative data of two independent experiments are shown. **(d)** Incubation time-dependent transduction of human fibroblasts by hFAP adapter-retargeted and untargeted Ad5. Recombinant Ad5 encoding TdTomato was pre-incubated with the hFAP adapter #6 and analyzed for transduction of hFAP^+^ D551 cells in comparison to the untargeted Ad5 after four, 24, and 48 hours incubation time of the (retargeted) adenoviral vector with the cells at an MOI of 10 (PFU/cell). Transduction was measured via cellular expression of TdTomato detected by flow cytometry. Representative data of two biological replicates of one experiment are shown.

### Generation and validation of DARPin-based hFAP adapters

To generate retargeting adapters for an hFAP-mediated cell transduction by Ad5, we used the selected hFAP-specific DARPins and incorporated them as retargeting module into our previously developed Ad5 adapter^22^. Protein analysis via SDS-PAGE and analytical SEC of the obtained adapters upon small-scale expression in *E. coli* and IMAC purification resulted in the selection of nine adapters. We subsequently tested these adapters for hFAP binding on (hFAP^+^) HT1080hFAP and (hFAP^−^) HT1080 cells. Our cell-based binding analysis showed specificity for all nine adapters, and found different adapter binding characteristics upon titration in a concentration range of 0.1 – 100 nM (Fig. 1c), confirming our previous data obtained with the single DARPins.

We hypothesized that not only the binding affinity but also the binding epitope, and/or other factors, play a role in the internalization process of the adenovirus into the cell. Therefore, we decided to test all of the selected adapters for their capability to promote an hFAP-dependent cell transduction.

To this end, we made use of both the HT1080hFAP and HT1080 cell line and performed cell transduction experiments with hFAP adapter-retargeted Ad5 (further on termed ‘hFAP-Ad5’) in comparison to the untargeted Ad5. Additionally, we included the non-targeting control adapter E3_5 with the Ad5 (further on termed ‘E3_5-Ad5’), which should block the fiber knob from interactions with cell surface molecules^22^ as it carries the non-binding DARPin E3_5. Cell transduction was determined via expression of the near-infrared fluorescent protein (iRFP) 670 encoded on the Ad5 vector and was detected via flow cytometry.

Six out of nine hFAP adapters (#3, #4, #5, #6, #8, #20) demonstrated a clear increase in the transduction of HT1080hFAP cells (60% – 90% transduced cells) when compared to the untargeted Ad5 (43% transduced cells; marked by the dashed line) (Fig. 1d). Out of these six retargeting adapters, two (#5, #8) also showed a major increase in transduction of hFAP^−^ HT1080 cells (20% – 45% transduced cells) when compared to the E3_5-Ad5 (9% transduced cells; marked by the dashed line), suggesting an unspecific, hFAP-independent transduction. Notably, these two adapters were both constituted of retargeting DARPins with two internal repeats (N2C), whereas the remaining adapters were constituted of retargeting DARPins with three internal repeats (N3C). As a result, we excluded these two adapters due to low specificity and selected four well-performing retargeting adapters (#3, #4, #6, #20) that enable a hFAP-mediated adenoviral cell transduction.

Interestingly, transduction levels observed with adapter #3 (constructed of a DARPin with comparably lower affinity) were equally high as those observed with adapters #4, #6, and #20 (all constructed of DARPins with comparably higher affinity), thus supporting our assumption that binding affinity alone does not determine a suitable Ad5 retargeting adapter.

Of further note, transduction levels of hFAP^−^ HT1080 cells were overall reduced in comparison to the transduction levels of hFAP^+^ HT1080hFAP cells. A possible explanation could be the cellular expression levels of the respective adenoviral attachment/entry molecule (CAR for untargeted Ad5 *versus* hFAP for hFAP-Ad5) depicted in Fig. 1b and Fig. 1e, thereby affirming our fiber knob adapter-based retargeting strategy.

### hFAP adapters enable Ad5 transduction of human fibroblasts

Having proven target specificity and functionality on the engineered fibrosarcoma cell line, we next aimed to validate the adapters for the retargeting of Ad5 via FAP in a more relevant setting using normal human fibroblasts. We chose the Detroit 551 (D551) human embryonic skin fibroblast cell line that endogenously expresses medium to high levels of hFAP (Fig. 2a) and low levels of the primary Ad5 attachment receptor CAR (Fig. 2b). Transduction of D551 cells by hFAP-Ad5, control E3_5-Ad5, or untargeted Ad5 revealed a massive increase (10 – 60 fold) in transduction when using the hFAP retargeting adapter for two different multiplicities of infection (MOIs; referring to plaque-forming units (PFU)/cell) (Fig. 2c). At an MOI of 20, all four hFAP adapter-retargeted Ad5 reached cell transduction levels of 42% to 58%, whereas the untargeted Ad5 showed only 5% cell transduction. The increased transduction efficiency mediated by the hFAP adapter was even more prominent at an MOI of two with transduction levels of 4% to 12% for all hFAP-Ad5 and only 0.5% for the untargeted Ad5. For the control E3_5-Ad5 we consistently observed a reduced transduction (10% at MOI 20; 0.7% at MOI 2) in comparison to all hFAP-Ad5, demonstrating that the increased transduction efficiency seen with the hFAP adapter was indeed mediated via the hFAP-specific retargeting DARPin.

Observing that the untargeted Ad5 only marginally transduced D551 cells, as opposed to the hFAP-Ad5, when incubated for six hours with the cells, we wondered whether the transduction levels would increase upon a longer incubation time of the vector with the cells. Therefore, we analyzed the transduction of D551 cells by hFAP adapter-retargeted and untargeted Ad5 upon 24 and 48 hours vector incubation time, as well as after only four hours vector incubation (Fig. 2d). For this experiment, we selected the so-far overall best-performing adapter #6 for Ad5 retargeting and chose an MOI of 10 (PFU/cell) based on our prior experiments. Furthermore, we used a TdTomato-encoding Ad5 since we found the signal intensity of this fluorescent protein to be stronger than the one detected with iRFP670 (data not shown). After four hours incubation time, the hFAP-Ad5 yielded 9.4% transduced cells, whereas the untargeted Ad5 yielded 0.3% transduced cells. For hFAP-Ad5, the transduction signal kept increasing over the time course, reaching 52.3% and 81.8% transduced cells after 24 and 48 hours incubation time, respectively. In contrast, the untargeted Ad5 yielded only 0.6% and 2.4% transduced cells after 24 and 48 hours incubation time, respectively, showing that even after a long duration the fibroblasts were rarely transduced by untargeted Ad5. Conversely, with the help of the hFAP adapter a considerable fraction of fibroblasts could be transduced by Ad5 after only a short duration. In line with the cellular hFAP and CAR expression levels (Fig. 2a and 2b), these data prove that the Ad5 transduction was truly hFAP-dependent and mediated via the hFAP retargeting adapter. Given that the human fibroblast D551 cell line is seemingly not susceptible to a natural Ad5 infection, our findings demonstrate the advantage of the hFAP adapter that enables Ad5 transduction of otherwise untransducable cells.

### Identification of a mFAP-specific adapter

Following our hFAP adapter screen and validation, we sought to generate an Ad5 adapter with specificity for murine FAP, enabling us to study vector retargeting to murine fibroblasts *in vivo* in mice. An initial sequence similarity search with the Basic Local Alignment Search Tool (BLAST; www.uniprot.org) revealed that the extracellular domains of human and murine FAP harbor 90.1% amino acid sequence identity, suggesting that our previously selected, hFAP-specific DARPins might be cross-reactive. To test this, we analyzed our well-behaved single DARPins (see Supplementary Fig. 1e) for binding to recombinant mFAP in an enzyme-linked immunosorbent assay (ELISA). Seven out of 18 hFAP DARPins indeed showed cross-reactivity to mFAP, as defined by a binding signal that was greater than the signal of our negative binding control (Fig. 3a). To confirm cross-reactivity and ensure mFAP-specificity on the cell surface, we analyzed binding using suitable target and non-target cells, as done before. Here, we used the mFAP-expressing, murine NIH3T3mFAP fibroblast cell line that had been created by lentiviral transduction of the FAP^−^ NIH3T3 parental cell line with murine FAP^40^ (Fig. 3b). Our cell-based binding analysis revealed that only candidate #6 of the seven presumably cross-reactive DARPins bound specifically to mFAP on cells, as shown by a high binding signal on target cells and no binding signal on non-target cells, while the remaining six DARPins consistently showed a higher binding signal on non-target than on target cells (Fig. 3c). This observation was confirmed with the corresponding DARPin-based adapters, which were as well tested for binding on NIH3T3mFAP and NIH3T3 cells (Supplementary Fig. 3). Therefore, we selected candidate #6 as both mFAP-specific and human/mouse cross-reactive adapter and proceeded to the *in vitro* Ad5 retargeting analysis.

**Figure 3:**
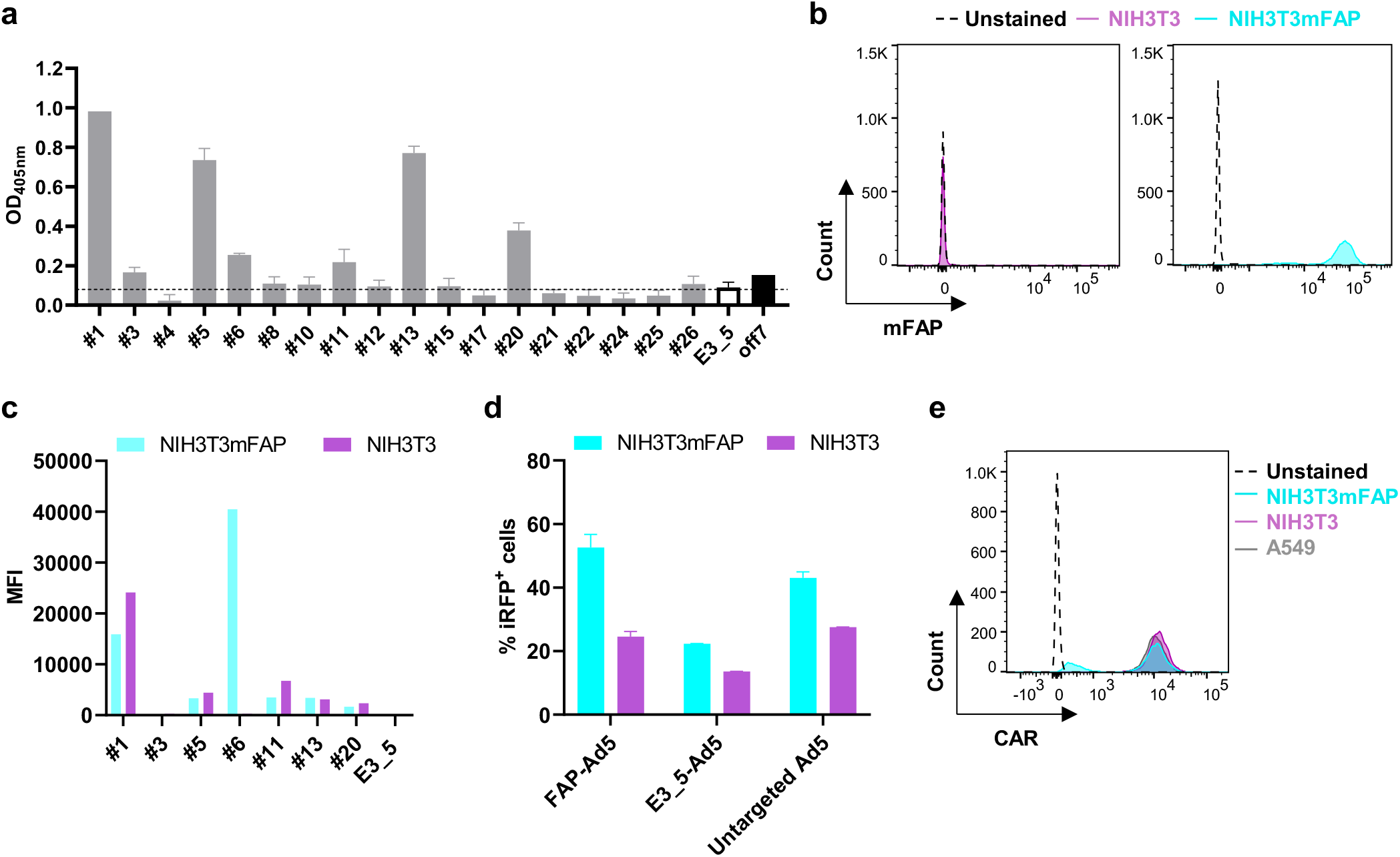
Retargeting of Ad5 to murine fibroblasts using a human/mouse cross-reactive FAP DARPin. **(a)** Analysis of hFAP DARPins for binding to mFAP via ELISA. Purified DARPins, previously selected to be hFAP-specific, were analyzed via ELISA for cross-reactivity to mFAP. The unselected, non-binding DARPin E3_5 was applied as a negative binding control. The maltose-binding protein (MBP)-specific DARPin off7 and recombinant MBP were applied as a technical positive binding control. The dashed line indicates a cut-off signal set on the negative binding control to select mFAP-binding DARPins. **(b)** Flow cytometry analysis of mFAP expression of the NIH3T3 and NIH3T3mFAP cell line with mFAP antibody staining. **(c)** Cell-based DARPin binding assay on target and non-target cells. Purified DARPins selected as mFAP binders by ELISA were analyzed for binding on mFAP^+^ NIH3T3mFAP and mFAP^-^ NIH3T3 cells. Binding was detected by flow cytometry upon FLAG-tag antibody staining of the FLAG-tagged DARPin. The unselected, non-binding DARPin E3_5 was applied as a negative binding control. Bars represent specific binding signal of single point measurements. Representative data at 1 µM DARPin concentration of a titration experiment are shown. MFI = Mean fluorescent intensity. **(d)** Transduction of target and non-target cells by mFAP adapter-retargeted Ad5. Recombinant Ad5 encoding iRFP670 was pre-incubated with the mFAP adapter #6 or the E3_5 blocking adapter and tested for transduction of mFAP^+^ NIH3T3mFAP and mFAP^−^ NIH3T3 cells in comparison to the untargeted Ad5 at an MOI of 2 (PFU/cell). Transduction levels were determined via cellular expression of iRFP670 detected by flow cytometry. Bars represent mean transduction level of two biological replicates ± SD. Representative data of three independent experiments are shown. **(e)** Flow cytometry analysis for CAR expression of the NIH3T3 and NIH3T3mFAP cell line in comparison to the positive control A549 cell line upon CAR antibody staining.

To test the selected mFAP-specific adapter for Ad5 retargeting to mFAP^+^ cells *in vitro*, we used an iRFP670-encoding Ad5 and analyzed transduction levels of NIH3T3mFAP and NIH3T3 cells upon incubation with mFAP-Ad5, untargeted Ad5, or control E3_5-Ad5 by flow cytometry. For NIH3T3mFAP cells, we detected a transduction rate of 52% for the mFAP-Ad5, 43% for the untargeted Ad5, and 22% for the control E3_5-Ad5 (Fig. 3d), demonstrating an increase in transduction when using the mFAP-specific adapter and a decrease in transduction when using the E3_5 blocking adapter. For NIH3T3 cells, we detected similar transduction levels for the mFAP-Ad5 and the untargeted Ad5 (25% and 28%, respectively) and a decreased transduction for the control E3_5-Ad5 (14%). Although the data verify the functionality of the mFAP adapter, a prominent retargeting effect as seen with the human fibroblast cell line (Fig. 2d) was not found with the murine cell line tested here. A possible explanation for this observation can be provided through the high CAR expression levels depicted in Fig. 3e, showing that both NIH3T3mFAP and NIH3T3 cells are highly positive for the primary Ad5 attachment receptor CAR. Consequently, high transduction levels can be achieved via both cell surface molecules, FAP and CAR, hampering a clear assessment of the contribution of the mFAP adapter. Nonetheless, since on mFAP^+^ cells we observed a decrease of transduction with the E3_5 control adapter (blocking CAR interaction) and an increase with the mFAP adapter compared to the untargeted Ad5, we conclude that the mFAP adapter is functional and promotes the retargeting of Ad5 to murine fibroblasts via FAP.

### Establishment of a FAP^+^ fibroblast-enriched tumor mouse model

Having selected and validated the mFAP-specific adapter in cell-based assays, we aimed to investigate its ability to mediate Ad5 retargeting to FAP^+^ cells *in vivo*. We chose the subcutaneous NCI-N87 human gastric cancer xenograft model utilizing immunodeficient SCID/beige mice. These mice are characterized by a lack of T and B lymphocytes, as well as by deficient natural killer (NK) cells, which facilitates tumor engraftment^46, 47^. NCI-N87 tumor cells overexpress HER2 but do not express FAP (Supplementary Fig. 4a). Therefore, vector retargeting to FAP^+^ stromal cells (and vector detargeting from tumor and other non-target cells) could be well monitored in this tumor mouse model.

**Figure 4:**
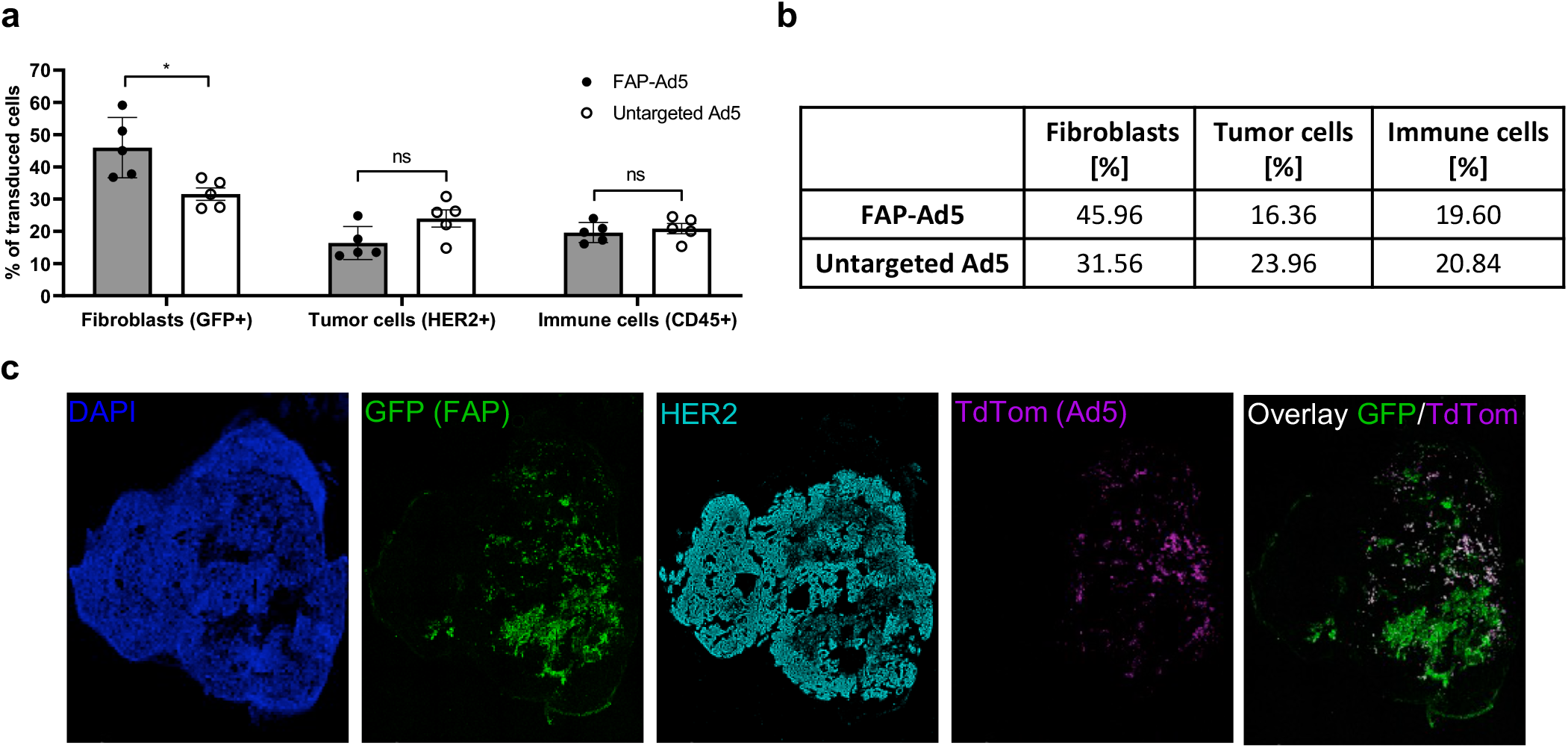
Successful retargeting of Ad5 to FAP-expressing fibroblasts *in vivo*. **(a)** HER2-overexpressing NCI-N87 tumor cells and GFP-labeled, mFAP-expressing NIH3T3mFAP fibroblast cells were co-injected subcutaneously into the flank of SCID/beige mice. After tumor establishment (200 mm^3^ tumor volume), mice were treated intratumorally with 3×10^9^ PFU FAP-retargeted or untargeted Ad5 encoding TdTomato. Three days post injection, tumors were harvested and analyzed by flow cytometry. Transduced cells were detected via TdTomato expression and further characterized by cell surface marker staining or GFP-expression. Each data point represents a single mouse. Bars represent mean ± SD of five mice per group. Statistics: Unpaired t-test; *p < 0.05. Representative data of two independent experiments are shown. **(b)** Quantification of (a), indicating mean values of transduced cells. **(c)** Immunohistochemical analysis of (a) to investigate the cell-specificity of FAP-retargeted Ad5. Representative immunofluorescence images of tumor tissues stained for HER2 (cyan) and counter-stained with DAPI (blue) for nuclei staining. FAP^+^ cells were detected via GFP-expression (green), and cells transduced with Ad5 were detected via TdTomato expression (magenta).

However, when we initially performed preliminary *in vivo* experiments to establish the tumor xenograft model, we found that the overall FAP-expression in the whole tumor was below 5% (Supplementary Fig. 4b). Reasonable modifications of the experimental protocol (e.g., number of tumor cells injected; time of tumor growth; age of the animal) did not significantly augment the FAP abundance. We observed the same limitation also for other subcutaneous tumor mouse models, including syngeneic models using immunocompetent C57BL/6 and BALB/c mouse strains. This is a major difference to human carcinomas which are commonly rich in CAFs^48^. Since a rather low abundance of FAP^+^ cells might impede a reliable Ad5 retargeting analysis, we thus established a FAP^+^ fibroblast-enriched subcutaneous tumor xenograft model, in which NCI-N87 tumor cells were co-injected with NIH3T3mFAP cells. These murine fibroblasts had been engineered to co-express mFAP^+^ and the green fluorescent protein (GFP), enabling their detection even without cell surface marker staining (Supplementary Fig. 4c). By varying the ratio of tumor cells to fibroblasts, this model then allowed an *in vivo* analysis of the cell-specificity as well as the cell-selectivity of the retargeted vector.

### Successful retargeting of Ad5 to FAP^+^ fibroblasts *in vivo*

To analyze mFAP adapter-mediated Ad5 retargeting *in vivo*, we co-injected NCI-N87 and NIH3T3mFAP cells subcutaneously into the flank of SCID/beige mice, using 100-fold more tumor cells than fibroblasts to not saturate the tumor with stromal cells and enable a fair retargeting analysis. After tumor establishment, and prior to administering the adenoviral vector, we examined the cellular composition of the tumor via flow cytometry (Supplementary Fig. 4d). On the day of vector injection, the tumor was comprised of 50% NIH3T3mFAP cells (GFP^+^), 30% tumor cells (HER2^+^), 5% immune cells (CD45^+^), and 15% other cells (unstained) (Supplementary Fig. 4e). Subsequently, we injected mFAP adapter-retargeted or untargeted TdTomato-encoding Ad5 intratumorally (i.t.), and analyzed cell transduction of harvested tumors via flow cytometry three days post injection.

We observed a significant increase in transduction of mFAP^+^ fibroblasts (46% FAP-Ad5 vs. 32% untargeted Ad5) and a considerable decrease in transduction of tumor cells (16% FAP-Ad5 vs. 24% untargeted Ad5) when applying the mFAP adapter to Ad5 (Fig. 4a and b), demonstrating a clear preference of the adapter-retargeted vector for targeted fibroblasts over off-target tumor cells. Regarding the transduction of immune cells, similar levels (20%) were detected for both FAP-Ad5 and untargeted Ad5.

To further investigate cell-specificity of the vector, we performed an immunofluorescence staining of the tumor tissues (Fig. 4c). Here, we again observed a strong preference of the FAP-retargeted Ad5 for mFAP^+^ fibroblasts, confirming our previous findings. As shown in the fluorescence microscopy images, the TdTomato signal of FAP-Ad5 predominantly co-localized with the GFP signal of mFAP^+^ fibroblasts. We therefore conclude that the adenoviral tropism could successfully be redirected to FAP^+^ stromal cells *in vivo* through the mFAP-specific DARPin adapter.

### Efficient *in vivo* delivery of active anti-cancer therapeutics using FAP-Ad5

We next explored whether the FAP-retargeted Ad5 would be functional for the *in vivo* delivery of cancer-attacking biomolecules to the TME. We used the FAP^+^ fibroblast-enriched subcutaneous NCI-N87 xenograft model in immunodeficient SCID/beige mice established here. This HER2-dependent human xenograft is sensitive to anti-HER2 therapy, allowing us to study a FAP-Ad5-mediated gene delivery of the mAb trastuzumab (TZB), for which an anti-tumor efficacy has previously been shown in this model^49^.

Accordingly, tumor-bearing mice received a single i.t. injection of FAP-retargeted TZB-encoding Ad5 (termed ‘FAP-Ad5-TZB’) and were then monitored for tumor growth for 19 days (Fig. 5a). For comparison, one group of tumor-bearing mice received a single i.t. injection of clinical-grade recombinant TZB (Herceptin^®^; termed ‘Herceptin 1x’), analogous to FAP-Ad5-TZB treatment, and another group received weekly-repeating i.t. injections of clinical-grade recombinant TZB for a total of three doses (termed ‘Herceptin 3x’). Tumor-bearing mice treated with PBS served as a control group.

**Figure 5:**
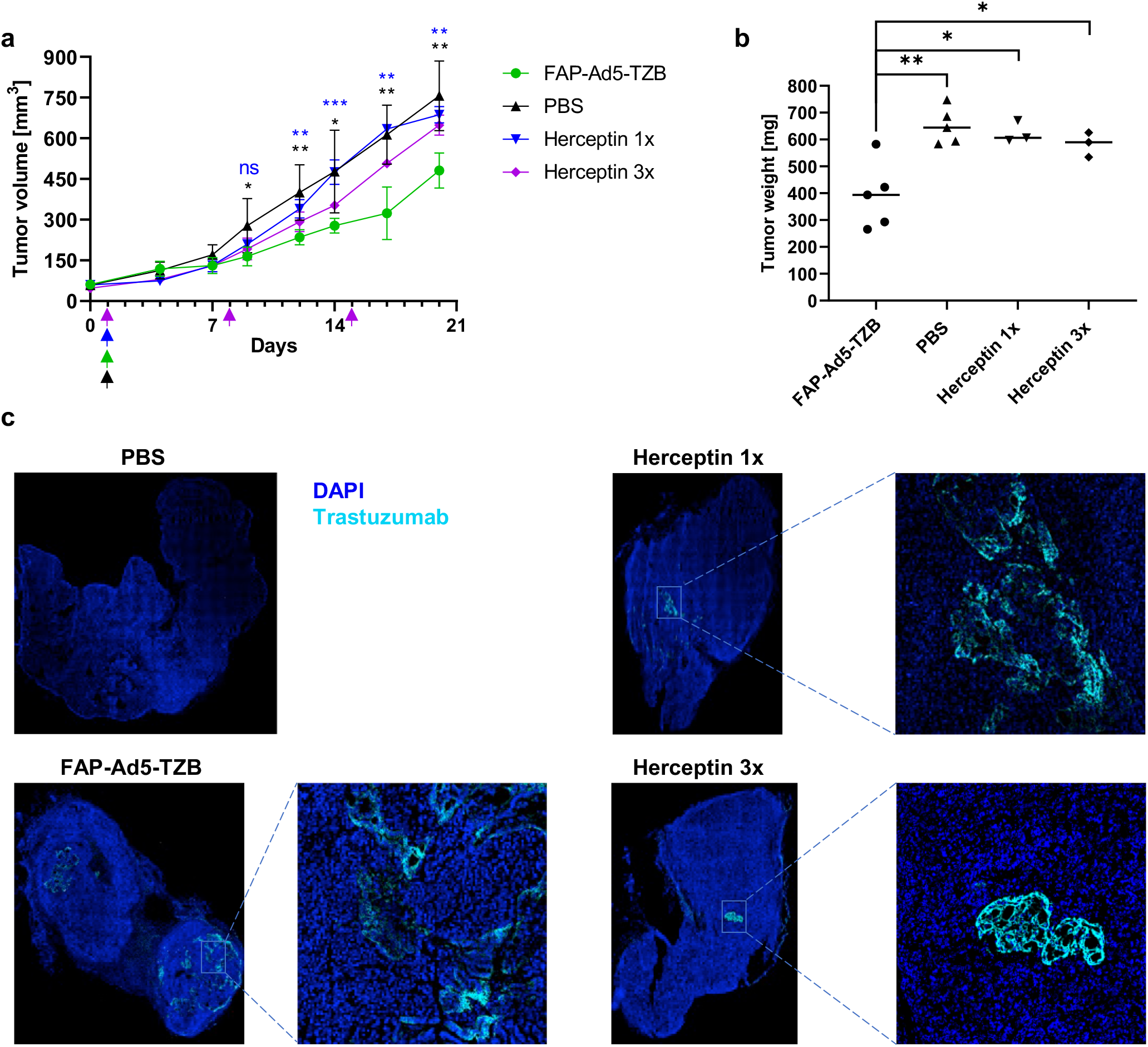
Efficient *in vivo* delivery of anti-cancer therapeutics using FAP-Ad5. **(a)** Growth analysis of HER2^+^ tumor xenografts upon Ad5-mediated treatment with trastuzumab (TZB) or with TZB as a protein. HER2-overexpressing NCI-N87 tumor cells and NIH3T3mFAP cells were co-injected subcutaneously into the flank of SCID/beige mice for tumor establishment. At a tumor volume of 50 mm^3^, mice were treated intratumorally with 9×10^8^ PFU FAP-retargeted Ad5 encoding trastuzumab (FAP-Ad5-TZB; n = 5), or one single dose of 200 µg Herceptin (n = 3), or three doses of 200 µg Herceptin (n = 3), or PBS (n = 5). Arrows indicate time points of injection for the corresponding treatment. Data points represent mean ± SD. Statistics: Unpaired t-test; *p < 0.05, **p < 0.005, ***p < 0.0005. **(b)** Tumor weights of harvested tumors from (a) 19 days post injection. Each data point represents a single mouse. Statistics: Unpaired t-test; *p < 0.05, **p < 0.005. **(c)** Detection of TZB within harvested tumors from (a) 19 days post injection. Representative immunofluorescence images of tumor tissues stained for TZB (cyan) and counter-stained with DAPI (blue) for nuclei staining.

We observed an immediate reduction in tumor growth throughout the first three days post injection for all Herceptin^®^-treated mice, whereas for FAP-Ad5-TZB-treated mice we observed the effect with a short delay throughout day four to seven (Fig. 5a). This short delay is presumably due to the initially required cellular expression of TZB after adenoviral transduction and the therefore delayed accumulation of the drug. Thereafter, tumors of all TZB-treated mice restarted growing, but interestingly much faster for Herceptin^®^-treated mice than for FAP-Ad5-TZB-treated mice, for which a significant delay in outgrowth could be detected. Tumor growth of Herceptin^®^-treated mice could moreover be slowed down through further injections of Herceptin (Herceptin 3x), leading us to the hypothesis that the profound delay in outgrowth seen for FAP-Ad5-TZB-treated mice is most likely attributed to a continuous production and secretion of the anti-cancer therapeutic throughout the study.

In line with these findings, tumors of FAP-Ad5-TZB-treated mice examined after tumor harvest were significantly smaller than those of PBS-or Herceptin^®^-treated mice (Fig. 5b and Supplementary Fig. 5).

To confirm the presence of TZB within the tumor, we analyzed tumor tissues via immunofluorescence staining to detect the therapeutic mAb. TZB could be detected in tumors of Herceptin^®^-treated as well as FAP-Ad5-TZB-treated mice but not in tumors of PBS-treated mice (Fig. 5c), correlating the presence of the drug with the observed biological effect. Remarkably, TZB was found to be present in various areas throughout the whole tumor for FAP-Ad5-TZB-treated mice, whereas for Herceptin^®^-treated mice TZB was found to be rather restricted to one spot within the tumor.

Taken together, our data demonstrate efficient *in vivo* delivery of a therapeutic payload to FAP^+^ fibroblasts in the TME by FAP-Ad5. The locally secreted biomolecule delayed tumor growth, thereby proving functionality of the FAP adapter to retarget Ad5 vectors to CAFs in the tumor stroma for cancer therapeutic applications.

## Discussion

Adenoviral vectors derived from Ad5 are one of the most widely used vectors for the delivery of gene therapeutics, and have shown great clinical promise in *ex vivo* and *in vivo* gene therapy^3^. However, *in vivo* gene delivery is still restricted due to the natural adenoviral tropism and insufficient vector targeting. In the present study, we developed a DARPin-based adapter with specificity for FAP to enable Ad5 retargeting to CAFs and thereby advance Ad5-mediated cancer gene therapy.

Using ribosome display together with a cell-based screening approach, we selected ten FAP-specific DARPins, partly with subnanomolar binding affinity, which served as targeting moiety in our modular adapter. Further selection and characterization of the eventually generated adapters yielded four candidates with specificity for hFAP. These four adapters enabled FAP-mediated and FAP-specific Ad5 transduction of FAP^+^ human fibroblasts *in vitro*, mimicking CAFs in the tumor stroma. Importantly, the FAP^+^ human fibroblasts expressed only low levels of the primary Ad5 cell attachment/entry receptor CAR and were thus hardly transduced by the untargeted Ad5, even after 48 hours of vector incubation. This experimental finding demonstrates the great advantage of our FAP retargeting adapter that enables gene delivery to FAP^+^ fibroblasts, and potentially other FAP^+^ cells, that are otherwise not transducable by Ad5 via CAR.

Amongst the final selected best-performing adapters, three showed exclusive binding to hFAP, whereas one showed binding to both hFAP and mFAP. This human/mouse cross-reactive adapter allowed us to perform preclinical experiments in mice, which can later be translated to patient-derived tumors. Our *in vivo* analysis of the cross-reactive FAP adapter demonstrated successful Ad5 retargeting to FAP^+^ fibroblasts in the TME in tumor-bearing mice. We observed a clear preference of the FAP-retargeted Ad5 for FAP^+^ fibroblasts and detected a decrease in FAP^−^ tumor cell transduction. Quantification of the cell populations present in the TME showed that at the time point of vector administration 50% of all cells consisted of target cells (fibroblasts), whereas the other 50% were composed of tumor cells, immune cells, and other non-target cells. The observed cell transduction within this heterogenous TME required a satisfactory cell-selectivity of the vector, which was indeed achieved through our retargeting module. Nonetheless, we did observe some residual off-targeting for FAP-Ad5 to the immune cell population, matching the transduction levels of the untargeted Ad5. We hypothesize that this off-targeting to immune cells mainly results from interactions of the hexon with macrophages, which are still abundant in the immunodeficient mouse model used here^50, 51^. Particularly HVR1 of the hexon is implicated in vector sequestration by macrophages through charge-dependent interactions with scavenger receptors located on these cells^50, 51^. In the present study, we did not apply the vector shield that has previously shown to mask HVR1 and other HVRs of the hexon^18^. The similar off-targeting to immune cells observed with both vectors could therefore indicate a hexon-dependent interaction promoting immune cell transduction, rather than a fiber knob-dependent interaction modifiable through our adapter molecule. Future studies using an HVR1-mutated vector, and/or applying our capsid shield, could be used to investigate our hypothesis further.

In contrast to our present study that comprises an *in vivo* retargeting analysis, Kuklik et al. recently presented an *in vitro* study on the retargeting of an AAV2-based vector to CAFs via FAP^52^. AAV2-based vectors equally rank amongst the most commonly used vectors for clinical gene therapy. However, AAV2 vectors harbor a relatively small packaging capacity (4.7 kb for AAV2 vs. 37 kb for Ad5), and are therefore restricted in payload size. Thus, high-capacity Ad5 vectors harbor a much larger packaging capacity and therefore allow for a delivery of large therapeutic gene cargos, with the opportunity of encoding regulatory elements and multiple genes. This important characteristic distinguishes our vector platform from other gene delivery approaches.

By using a HER2-overexpressing mouse tumor xenograft model, we could furthermore test *in vivo* gene delivery of the clinically relevant mAb TZB to FAP^+^ stromal fibroblasts by our FAP-retargeted Ad5. FAP-Ad5-delivered TZB showed superior efficacy over clinical-grade recombinant TZB (Herceptin^®^) applied as a protein via the same route of administration. We attribute this superiority of FAP-Ad5-delivered TZB to a presumably continuous production and secretion of the drug by transduced fibroblasts throughout the study, leading to a more even biodistribution within the TME. We had previously studied the production of TZB by the tumor cells themselves, upon delivery via a HER2-retargeted Ad5 to HER2^+^ tumor cells, and observed a long duration of production^53^. Indeed, TZB had been found in the TME 61 days post i.t. vector injection, indicating a continuous secretion of the therapeutic mAb, which shows a half-life of 28 days in humans but of only eight days in mice^54, 55^. The sustained biomolecule expression constitutes another benefit of our adenoviral delivery platform in comparison to a direct drug administration, particularly for molecules with short half-life. This characteristic and the concept of a local drug production to improve bioavailability and reduce off-target toxicities reaffirmed our aim of installing a ‘biofactory’ in the TME.

To this end, the stromal cell-targeted (paracrine) delivery approach presented here might exhibit certain advantages over our previous tumor cell-targeted (autocrine) delivery: (i) expression of the anti-cancer agent in the TME might be of longer duration in the paracrine approach since the ‘biofactory’ (transduced CAFs) would not be therapeutically affected by the encoded agent. In contrast, in an autocrine approach, expression of the anti-cancer agent might be limited by the lifetime of the transduced (tumor) cells, as they are producer cells but at the same time affected by the action of the therapeutic. (ii) Targeting the vector to CAFs might also be beneficial regarding the spatial distribution and/or abundance of stromal cells, particularly in stroma-rich tumors which are common for human cancers^48^. (iii) Given that FAP is expressed in more than 90% of all carcinomas^28^, a stromal-cell targeted delivery via FAP on CAFs would moreover allow to treat a variety of human cancers, independent of the existence of specific/targetable tumor cell markers or suitable retargeting adapters. This might additionally be a promising strategy to avoid antigen escape of even those tumors cells that carry defined markers.

In conclusion, the FAP-specific adapter presented herein expands our Ad5 retargeting system, and enables a stromal cell-targeted delivery of anti-cancer biomolecules to the TME with potential benefits over existing, tumor cell-targeted delivery strategies. The FAP adapter can be combined with our scFv-based capsid shield^18^ to overcome pre-existing immunity against the vector^20^. More importantly, it can be applied to any Ad5-derived vector including high-capacity Ad5 vectors, which allows the simultaneous delivery of multiple payloads for combination cancer therapy as shown by Brücher et al.^56^. Thus, our FAP-retargeted Ad5 delivery platform may help advance current cancer treatment options involving biologics with required local or prolonged production. Beyond cancer therapy, our FAP-Ad5 vector could potentially be applied to other medical disorders with FAP-involvement, such as chronic inflammatory diseases and fibrosis^57-61^.

## Methods

### Chemicals

All chemicals were purchased from Sigma-Aldrich unless stated otherwise.

### Cell lines

The cell lines HT1080 and HT1080hFAP were generated as described previously^39^ and kindly provided by Prof. Christian Münz (University of Zürich). HT1080 and HT1080hFAP cells were cultured in complete RPMI media (RPMI1640 GlutaMax (Thermo Fisher Scientific) supplemented with 10% (v/v) fetal bovine serum (FBS; BioConcept), 100 U/ml penicillin and 100 μg/mL streptomycin (Sigma-Aldrich)) containing 200 μg/mL G418 (Carl Roth) and 150 μg/mL Hygromycin B (Invitrogen). NIH3T3 and NIH3T3mFAP cell lines were generated as described previously^40^ and kindly provided by Prof. Ellen Puré (University of Pennsylvania). NIH3T3 and NIH3T3mFAP cells were cultured in complete RPMI media. HEK293, D551, HeLa, A549, and NCI-N87 cell lines were purchased from the American Type Culture Collection (ATCC). HEK293, D551, HeLa, and A549 cells were cultured in complete DMEM media (DMEM-high glucose (Sigma-Aldrich) supplemented with 10% (v/v) FBS, 100 U/ml penicillin and 100 μg/mL streptomycin), and NCI-N87 cells were cultured in complete RPMI media. All cell lines were maintained at 37 °C and 5% CO_2_ in a humidified atmosphere, and routinely tested and confirmed negative for mycoplasma contamination.

### Ribosome display selection of DARPins against hFAP

Ribosome display selections were performed as described previously^44^ using a semi-automatic KingFisher Flex MTP96 well platform. Biotinylated recombinant human FAP (hFAP; Sino Biological) was used for the selection, in alternating selection rounds, on either MyOne T1 streptavidin-coated beads (Pierce) or Sera-Mag neutravidin-coated beads (GE Healthcare), and a total of four selection rounds were performed. After *E. coli* transformation, binders were directly analyzed for hFAP binding by performing flow cytometry-based cell-binding assays described below.

### Cloning, expression, and purification of DARPins and adapters

DARPins were cloned into the pQIq backbone containing an N-terminal His_6_-tag and a C-terminal FLAG-tag. Adapters were cloned into the pQIq backbone containing an N-terminal His_10_-tag with a 3C protease cleavage site. For DNA propagation, the *E. coli* XL1-Blue strain was transformed with the corresponding plasmids.

For high-throughput screening, DARPins and adapters were expressed in a small-scale format (1 mL culture volume, 96-deep-well plate (Thermo Fisher Scientific)) in the *E. coli* XL1-Blue strain. Upon cell lysis, proteins were purified via immobilized-metal ion affinity chromatography (IMAC) using a HisPur™ Cobalt Spin Plate (Thermo Fisher Scientific) equilibrated with equilibration buffer (50 mM Na_2_HPO_4_/NaH_2_PO_4_, 300 mM NaCl, pH 7.4). After washing with wash buffer (50 mM Na_2_HPO_4_/NaH_2_PO_4_, 300 mM NaCl, 15 mM imidazole, pH 7.4), proteins were eluted in elution buffer (50 mM Na_2_HPO_4_/NaH_2_PO_4_, 300 mM NaCl, 250 mM imidazole, pH 7.4) and further buffer-exchanged to PBS (pH 7.4) using a Pall AcroPrep™ filter plate (Pall). Purity and molecular weights were analyzed via sodium dodecyl sulfate polyacrylamide gel electrophoresis (SDS-PAGE) and analytical size exclusion chromatography (SEC) using a Superdex™ 75 5/150 GL column (GE Healthcare).

For *in vivo* experiments, selected adapters were expressed in large-scale format (500 mL culture volume) in the *E. coli* BL21 strain. Upon cell lysis, protein purification was carried out via IMAC using Ni-NTA agarose (Qiagen) equilibrated with TBS_Eq_ (50 mM Tris-HCl, 400 mM NaCl, pH 7.4). After washing with TBS_W_ (50 mM Tris-HCl, 400 mM NaCl, 20 mM imidazole, 10% (v/v) glycerol, pH 7.4), adapters were eluted in TBS_E_ (50 mM Tris-HCl, 400 mM NaCl, 250 mM imidazole, pH 7.4). Purified adapters were then incubated with 3C protease (kindly provided by Dr. Fabian Brandl, University of Zurich) for removal of the His_10_-tag and dialyzed against PBS (pH 7.4) for buffer exchange. Adapters were then further purified via preparative SEC using a Superdex™ 200 10/300 GL column (GE Healthcare). Purity and molecular weights were analyzed as described above. Prior to *in vivo* injection into mice, purified adapters were assessed for endotoxin content using the EndoSafe® Portable Test System™ (Charles River Laboratories) and test cartridges with 0.5 – 0.005 EU/mL sensitivity, to confirm that endotoxin levels would not exceed the limit for endotoxin content for injectable products recommended by the Food and Drug Administration (FDA).

### Flow cytometry-based cell-binding assay for binder screening

For a high-throughput target-binding analysis of DARPins and adapters, we performed flow cytometry-based cell-binding assays using suitable target and non-target cells as follows. Cells were harvested by trypsinization and washed with ice-cold PBS-B (DPBS (Sigma-Aldrich) containing 1% (w/v) bovine serum albumin (BSA). 1×10^5^ cells resuspended in PBS-B were added per well of a 96-well plate (V-bottom, non-treated surface; Thermo Fisher Scientific) and kept on ice. Purified DARPins or adapters diluted in DPBS (or crude *E. coli* extracts in a final dilution of 1:100) were added to the cells and incubated for 60 minutes (min) on ice. Thereafter, cells were washed with ice-cold PBS-B and subsequently stained for 30 min on ice with appropriate antibodies diluted in PBS-B to detect the bound DARPin or adapter. Mouse anti-FLAG M2-FITC antibody (Sigma-Aldrich, F4049; 1:200) was used for DARPin detection. Mouse anti-His (Qiagen, 34670; 1:200) and goat anti-mouse IgG-AF488 (Invitrogen, A11001; 1:1000) were used for adapter detection. Upon antibody staining, cells were washed with ice-cold PBS-B and then fixed with fixation buffer (DBPS containing 2% (w/v) paraformaldehyde (PFA) for 10 min at room temperature (RT).

After a final wash step, cells were resuspended in PBS-B. Samples were measured on a BD FACSCanto™ II (BD Biosciences) using the high-throughput sampler, and flow cytometry analysis was performed using FlowJo™ software (BD Biosciences).

### Enzyme-linked immunosorbent assay (ELISA)

Mouse FAP (mFAP)-binding DARPins were screened via ELISA using recombinant mFAP (R&D Systems), and recombinant maltose-binding protein (MBP; kindly provided by Joana Marinho, University of Zurich) for an MBP binder as a positive control. Target protein was coated overnight (O/N) at 4 °C on MaxiSorp™ 96-well plates (Nunc). ELISA-blocking buffer (PBS containing 0.5% (w/v) BSA) was used for blocking performed for 2 h at RT while shaking on a plate shaker. Plates were washed with ELISA-PBS-T (PBS containing 0.05% (v/v) Tween-20). Samples were diluted (100 nM DARPin concentration) in ELISA-PBS-TB (ELISA-PBS-T containing 0.5% (w/v) BSA) and incubated for 1 h at 4 °C while shaking. DARPins were detected with rabbit anti-FLAG antibody (GenScript, A01868; 1:5,000) and goat anti-rabbit-AP antibody (Sigma-Aldrich, A3687; 1:10,000), both diluted in ELISA-PBS-TB and incubated for 1 h at 4 °C while shaking. Substrate solution consisting of 3 mM p-nitrophenyl phosphate (pNPP) diluted in pNPP-buffer (50 mM NaHCO_3_, 50 mM MgCl_2_) was used for final detection. Absorbance at 405 nm was measured at Tecan Infinite M1000 plate reader (Tecan).

### Affinity measurements using surface plasmon resonance (SPR)

Binding kinetics of selected DARPins were determined as previously described^62^ using recombinant biotinylated hFAP (Sino Biological). In brief, recombinant biotinylated hFAP was diluted in PBS (pH 7.4) and immobilized on a SPPNAHC200M Sensor Chip (XanTec Bioanalytics) to a level of 1000 resonance units. DARPins were diluted in running buffer (PBS (pH 7.4) containing 0.005% (v/v) Tween-20) covering a concentration range of 0.1 to 10 nM. A 1:1 Langmuir binding model was used to fit the data measured on a ProteOn™ XPR36 instrument (Bio-Rad Laboratories) using ProteOn™ Manager Software (Version 3.1.0.6, Bio-Rad Laboratories).

### Viral vector generation and purification

All Ad5 vectors used in this study were E1/E3-deleted and thus replication-deficient. Furthermore, all Ad5 vectors harbored four genetic mutations in the hexon hypervariable region 7 (I421G, T423N, E424S and L426Y) to reduce coagulation factor X-mediated liver tropism, as described previously by Schmid et al.^18^. Vectors were generated as described before^18, 53^ using the AdEasy Adenoviral Vector System (Agilent). Vector purification was performed via ultracentrifugation using two sequential cesium chloride gradients. In some instances, generated vectors were reamplified and purified by Vector Biolabs (Malvern, PA, USA).

### Flow cytometry analysis

Cell marker expression was determined via flow cytometry upon antibody staining as follows. For cell lines, cells were harvested by trypsinization, washed with ice-cold FACS-buffer (DPBS containing 1% (w/v) BSA and 0.1% (w/v) NaN_3_), and resuspended in ice-cold FACS-buffer to proceed with the antibody staining. Antibody staining was performed by incubating 1×10^5^ – 1×10^6^ cells with the appropriate antibody (see list below) for 20 min on ice. Cells were then washed with ice-cold FACS-buffer and fixed with fixation buffer for 10 min at RT. After a final wash step, cells were resuspended in FACS-buffer and stored at 4 °C until being analyzed at the flow cytometer.

For harvested solid tumors, tumors were minced with a scalpel, incubated in digestion media (RPMI1640 GlutaMax supplemented with 2% (v/v) FBS, 1 mg/mL Collagenase IV (Gibco), 0.5 mg/mL Hyaluronidase (ITW Reagents), and 0.5 mg/mL DNase I (Merck)) for 30 min at 37 °C, and passed through a 70 µm mesh to yield a single-cell suspension. Cells were collected by centrifugation (500×g for 5 min at 4 °C) and washed with ice-cold DPBS. Thereafter, red blood cell (RBC) lysis was performed using ACK Lysing Buffer (Thermo Fisher Scientific) following the manufacturer’s instructions. ACK Lysing Buffer was diluted by adding a tenfold excess of DPBS containing 5 mM EDTA, followed by an incubation for 10 min on ice. Subsequently, cells were washed with ice-cold DPBS and subjected to a live/dead staining using the LIVE/DEAD™ Fixable Violet Dead Cell Stain Kit (Invitrogen). Then, IgG Fc receptors were blocked using TruStainFcX™ PLUS (BioLegend) followed by the antibody staining procedure described above.

For multicolor flow cytometry panels, single-stained controls or UltraComp eBeads Plus (Thermo Fisher Scientific) were used for compensation. Fluorescence minus one (FMO) and further proper controls were included in each experiment.

Flow cytometers used in this study included BD FACSCanto™ II, BD LSRFortessa™, BD FACSymphony™ 5L (all BD Biosciences), and Cytek Aurora 5L (Cytek Biosciences). Flow cytometry analysis was performed using FlowJo™ software (BD Biosciences).

### Antibodies used for flow cytometry

Antibodies used for flow cytometry included: mouse anti-human FAP-APC (R&D Systems, FAB3715A), sheep anti-human FAP (R&D Systems, AF3715), mouse anti-mouse FAP (Merck, MABC1145), mouse anti-CAR (Millipore, 05-644), mouse anti-HER2-AF488 (BioLegend, 324410), mouse anti-HER2-AF647 (BioLegend, 324412), rat anti-mouse CD45-AF700 (BioLegend, 103127), donkey anti-sheep IgG-CF488A (Sigma-Aldrich, SAB4600038), goat anti-mouse IgG-AF647 (Invitrogen, A21235), goat anti-mouse IgG-AF488 (Invitrogen, A11001). Optimal antibody dilutions were determined via titration or adopted from the manufacturer’s recommendation.

### *In vitro* Ad5 transduction assay

1×10^4^ or 5×10^4^ cells were seeded per well of a cell culture 96-well or 24-well plate (Corning), respectively. The Ad5 vector was pre-incubated with the corresponding adapter (tenfold molar excess of adapter over knob) diluted in DPBS for 1 hour (h) on ice. Untargeted vector was treated accordingly by substituting the adapter by DPBS only. For transduction, cell medium was changed to fresh complete media and (retargeted) Ad5 vector was added to the cells at the indicated multiplicity of infection (MOI; referring to plaque-forming units (PFU)/cell). Cells were incubated for 6 h (unless stated otherwise) at 37 °C and 5% CO_2_ before performing another medium change to remove the viral vector. Cells were further incubated for 20 – 26 h at 37 °C and 5% CO_2_ and then harvested by trypsinization. If needed, antibody staining to detect cell surface markers was performed as described above. After washing, cells were fixed with fixation buffer for 10 min at RT and thereafter stored in FACS-buffer at 4 °C until being analyzed at the flow cytometer (instruments listed above). Transduction was measured via the cellular expression of a fluorescent reporter protein (iRFP670 or TdTomato) for which the corresponding DNA had been delivered by the Ad5 vector.

### Animal experiments

Eight-to-nine weeks old female Fox Chase SCID/beige mice (CB17.Cg-Prkdc^scid^Lyst^bg-J^/Crl; Charles River) were used for the *in vivo* studies. All animal experiments were performed in accordance with the Swiss animal protection law and with approval of the Cantonal Veterinary Office (Zurich, Switzerland).

### *In vivo* Ad5 retargeting study

Mice were subcutaneously injected into the left flank with 1×10^6^ NCI-N87 and 1×10^4^ NIH3T3mFAP cells in 100 µL DPBS containing 50% Matrigel (Corning). Tumors were measured with a caliper and tumor volumes calculated from V = 0.5 × length × width × width. Nineteen days after tumor engraftment (reaching approximately 200 mm^3^ tumor volume), mice were intratumorally injected with 50 µL of indicated Ad5 vector (3×10^9^ PFU in DPBS) encoding TdTomato. For FAP-retargeted Ad5, vector had been pre-incubated with the corresponding adapter (tenfold molar excess of adapter over knob) diluted in DPBS for 1 h on ice. Three days post injection, mice were sacrificed, and tumors were harvested and processed for subsequent analysis via flow cytometry (see above) or immunohistochemistry (see below).

### *In vivo* Ad5 therapeutic delivery study

Mice were subcutaneously injected into the left flank with 1×10^6^ NCI-N87 and 1×10^5^ NIH3T3mFAP cells in 100 µL DPBS containing 50% Matrigel. Tumor volumes were determined as stated above. Nine days after tumor engraftment (reaching approximately 50 mm^3^ tumor volume), mice were intratumorally injected with 50 µL of either i) FAP-Ad5-TZB (FAP-retargeted Ad5 encoding trastuzumab; for Ad5 retargeting procedure see above; 9×10^8^ PFU in DPBS), or ii) Herceptin^®^ (Roche; 200 µg in DPBS), or iii) PBS (DPBS). One group of mice received repeated intratumoral injections (one injection per week) of Herceptin^®^ (200 µg in DPBS) adding up to a total of three injections. Nineteen days post (first) injection, mice were sacrificed and tumors were harvested, weighted, and processed for subsequent analysis via immunohistochemistry (see below).

### Immunohistochemistry (IHC)

IHC was performed using cryosections (10 μm thick) of frozen tumor tissues embedded in O.C.T.™ compound (Thermo Fisher Scientific). Cryosections were fixed with acetone for 10 min at −20 °C, washed with IHC-PBS-T (PBS (pH 7.4) containing 0.1% (v/v) Tween-20), and blocked with IHC-blocking buffer (IHC-PBS-T with 10% normal goat serum (Cell Signaling Technology)) for 1 h at RT. Sections were incubated with primary antibody diluted in IHC-blocking buffer O/N at 4 °C. After washing, sections were incubated with appropriate secondary antibodies diluted in IHC-blocking buffer for 1 h at 4 °C, washed, and counterstained with DAPI (Thermo Fisher Scientific; 300 nM final concentration) for 5 min at RT. After final washing, sections were mounted with ProLong Gold antifade mountant (Thermo Fisher Scientific) and analyzed using the THUNDER Imager Microscope (Leica). Antibodies used for IHC analysis included rat anti-HER2 (Thermo Fisher Scientific, MA1-82367; 1:500), goat anti-rat IgG-AF647 (Amersham Biosciences, 4418; 1:1000), and goat anti-human IgG-AF647 (Thermo Fisher Scientific, A21445; 1:1000).

## Acknowledgement

We are grateful to Joana Marinho, Thomas Reinberg, Anna-Lena Schinke, and Sven Furler for DARPin selection and hit validation, and to Dr. Jonas Schäfer and Dr. Birgit Dreier for supervising the activity. We thank Prof. Ellen Puré and Prof. Christian Münz for providing cell lines, Dr. Lukas Becker for helping with protein purification and for scientific discussions, Nina Schumacher and Faye Menard for helping with tumor processing during *in vivo* studies, Dr. Fabian Brandl for assisting with SPR analysis and for providing reagents, Dr. Markus Schmid for providing materials and reagents, and Jonas Kolibius for scientific discussions and critical reading of the manuscript. We are grateful to the Zurich Integrative Rodent Physiology (ZIRP) team, especially to Dr. Svende Pfundstein for training and technical assistance during *in vivo* experiments, as well as to Dr. Petra Seebeck for assistance with the animal license application. We acknowledge the Flow Cytometry Facility of the University of Zurich for training and maintenance of the instruments. We further acknowledge the Laboratory Animal Services Center of the University of Zurich for training, animal caretaking, and experimental equipment.

## Author contributions

K.P.H.: Conceptualization, Methodology, Investigation, Validation, Formal analysis, Visualization, Project administration, Writing - Original Draft;

M.v.G.: Investigation, Resources, Writing - Review & Editing;

P.C.F.: Resources, Visualization, Writing - Review & Editing;

F.K.: Methodology, Writing - Review & Editing;

G.N.-D.: Investigation, Writing - Review & Editing;

L.B.: Resources, Writing - Review & Editing;

A.P.: Supervision, Conceptualization, Funding acquisition, Writing - Review & Editing;

