## Supplementary Figures for "FAP-retargeted Ad5 enables *in vivo* gene delivery to stromal cells in the tumor microenvironment"

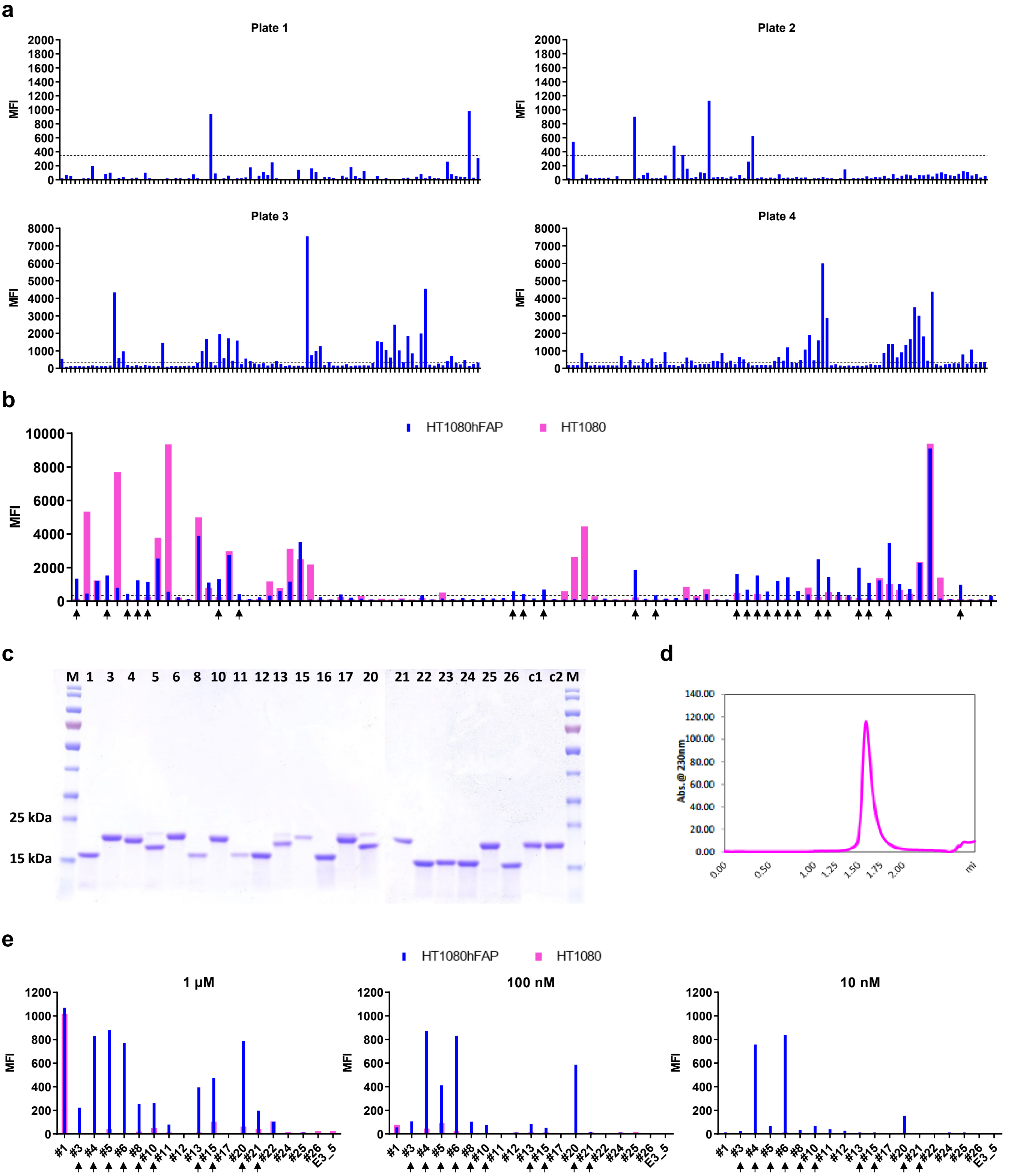


**Supplementary Figure 1: Selection of DARPins against hFAP.** Crude extract cell-binding assay on **(a)** target and **(b)** target and non-target cells. Crude extracts of DARPin-expressing *E. coli* were tested for binding on hFAP$^{+}$ HT1080hFAP cells (blue bars) and hFAP$^{-}$ HT1080 cells (pink bars) at 1:100 dilution. Binding was detected via flow cytometry by staining of the FLAG-tagged DARPin. The unselected DARPin E3_5 was applied as a non-binding control yielding a cut-off signal (dashed line) to identify hFAP-binding

DARPins. Arrows in (b) indicate the ‘Top 25‘ hFAP-specific DARPins selected upon side-by-side binding comparison on target and non-target cells. Bars represent binding signal of single-point measurements. Representative data of three independent experiments are shown. MFI = Mean fluorescent intensity. **(c)** SDS-PAGE analysis of purified ‘Top 20‘ hFAP DARPins. The selected ‘Top 20‘ hFAP-binding DARPins and two control DARPins (the MBP-specific DARPin off7 denoted as ‘c1‘, and the unselected DARPin E3_5 denoted as ‘c2‘) were expressed in *E. coli* and purified via their His-tag by IMAC. Purified proteins were analyzed on a 12% SDS-polyacrylamide gel. DARPins with two (N2C) or three (N3C) internal repeats are expected at a molecular weight of 16 kDa or 19 kDa, respectively. It should be noted that some DARPins do not unfold under these conditions and thus run as more compact proteins. M = molecular weight marker; kDa = kilodalton. **(d)** SEC analysis of purified ‘Top 20‘ hFAP DARPins. Purified hFAP DARPins were analyzed by gel filtration to identify monodisperse proteins with an elution profile corresponding to the representative graph shown here. **(e)** DARPin cell-binding assay on target and non-target cells. Using three different concentrations, purified hFAP DARPins were analyzed in parallel for binding on $hFAP^+$ HT1080hFAP cells (blue bars) and $hFAP^-$ HT1080 cells (pink bars). Binding was detected via flow cytometry by staining of the FLAG-tagged DARPin. The unselected E3_5 DARPin was applied as a non-binding control. Arrows indicate the ‘Top 10‘ purified DARPins specific for hFAP.

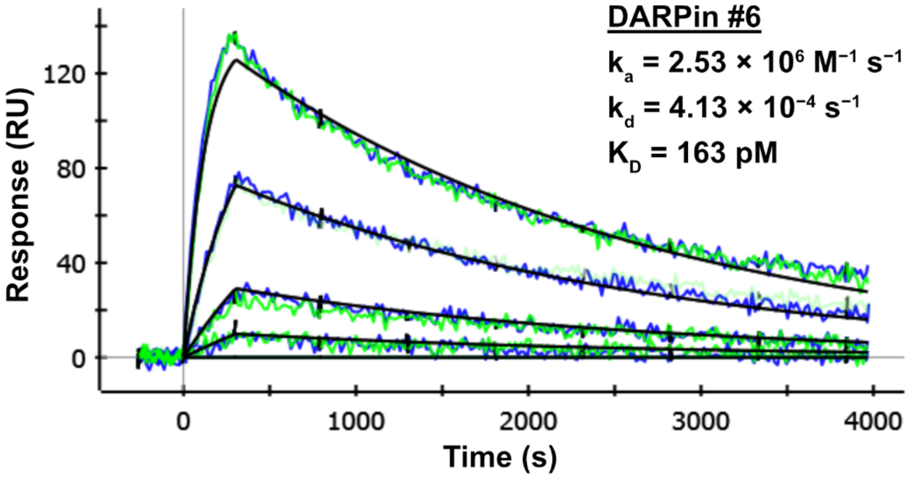


**Supplementary Figure 2: Affinity determination of hFAP-binding DARPin #6.** SPR experiments were performed using immobilized biotinylated recombinant hFAP and different dilutions of DARPin #6 covering a concentration range of 0.1 to 10 nM. Blue and green curves represent duplicate measurements whereas the black curve represents the respective fit.

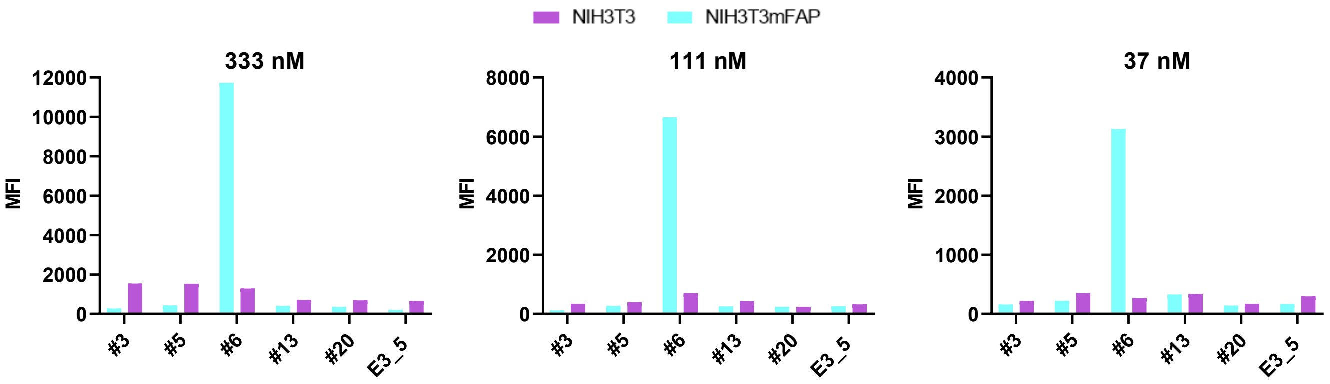


**Supplementary Figure 3: Selection of a mFAP-specific adapter to retarget Ad5 to murine fibroblasts.** Cell-based adapter binding assays on target and non-target cells. Selected purified adapters were analyzed for binding on mFAP$^{-}$ NIH3T3 and mFAP$^{+}$ NIH3T3mFAP cells at three different concentrations. Binding was detected via flow cytometry by staining of the His-tagged adapter. The unselected, non-binding adapter E3_5 was applied as a negative binding control. Bars represent binding signal of single-point measurements. MFI = Mean fluorescent intensity.

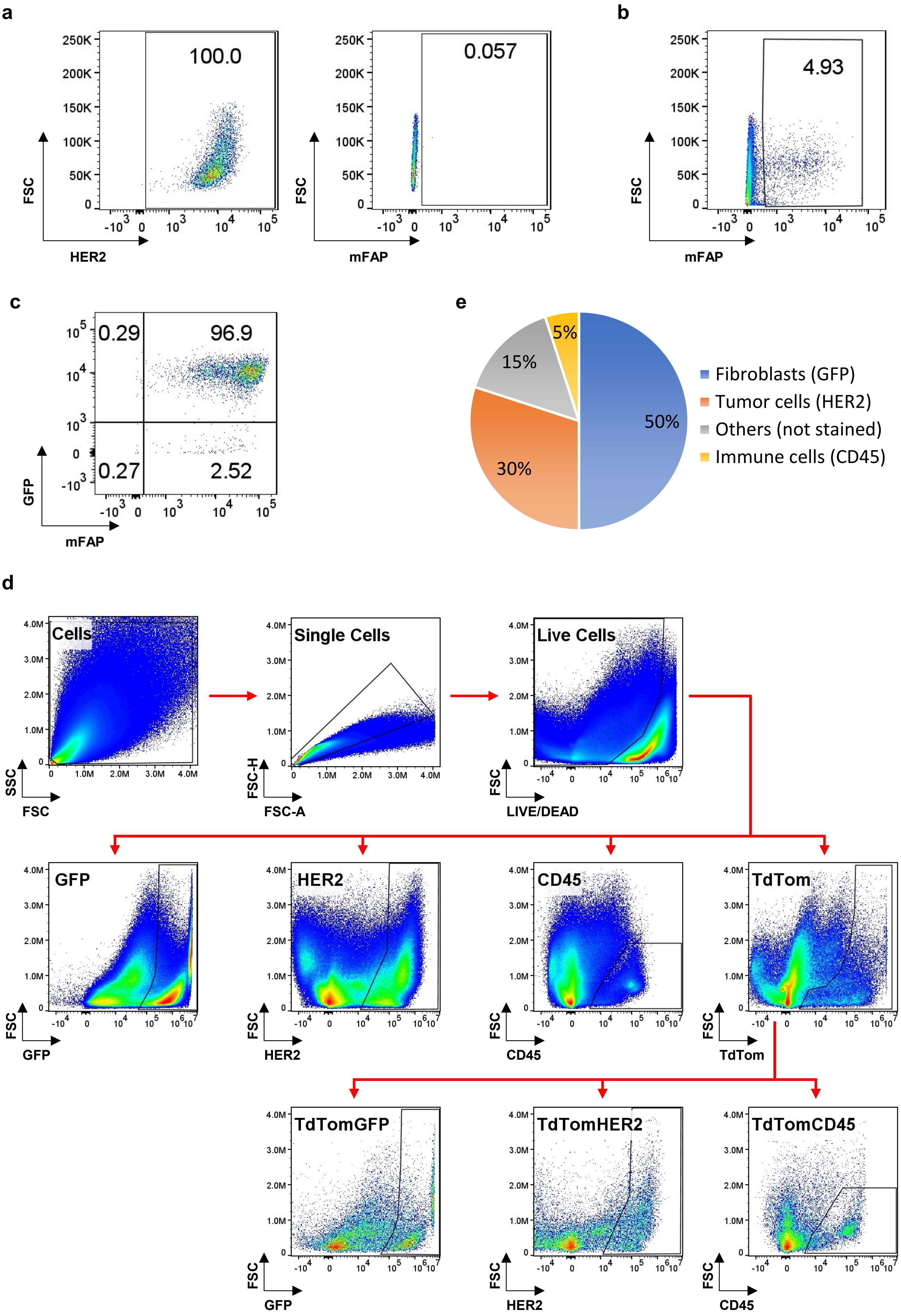

a
100.0
0.057
HER2
mFAP
FSC
b
4.93
c
0.29
96.9
0.27
2.52
GFP
d
Cells
Single Cells
Live Cells
SSC
FSC
FSC-H
FSC-A
LIVE/DEAD
GFP
HER2
CD45
TdTom
TdTomGFP
TdTomHER2
TdTomCD45
e
50%
30%
15%
5%
Fibroblasts (GFP)
Tumor cells (HER2)
Others (not stained)
Immune cells (CD45)

**Supplementary Figure 4: Establishment and characterization of a FAP$^+$ fibroblast-enriched mouse tumor model.** **(a)** Flow cytometry analysis for HER2 and mFAP expression of the NCI-N87 human gastric cancer cell line. **(b)** Flow cytometry analysis for mFAP expression of a subcutaneous NCI-N87 tumor xenograft. NCI-N87 tumor cells were injected subcutaneously into the flank of SCID/beige mice, established for three weeks, and harvested at a tumor volume of 400 $mm^3$ to determine mFAP expression. **(c)** Flow cytometry analysis of NIH3T3mFAP cells for GFP and mFAP expression. **(d)** Gating strategy applied for the tumor flow cytometry analysis to determine the cellular composition and to detect Ad5-transduced cells. Cells were stained with LIVE/DEAD$^{TM}$ Fixable Violet Stain Kit and fluorescently-labeled HER2- and CD45-specific antibodies. To determine the cellular composition of the tumor, live single cells were then gated for GFP or HER2 or CD45 to quantify fibroblasts, tumor cells or immune cells, respectively. To quantify the amount of Ad5-transduced cells, live single cells were first gated for TdTomato and then for GFP or HER2 or CD45 to quantify transduced fibroblasts, tumor cells or immune cells, respectively. **(e)** Cellular composition of the FAP$^+$ fibroblast-enriched tumor on the injection day of FAP-retargeted or untargeted Ad5 during the *in vivo* retargeting study. NCI-N87 tumor cells and NIH3T3mFAP cells were co-injected subcutaneously into the flank of SCID/beige mice. On the day of adenoviral vector injection, two mice of the control group were sacrificed for tumor harvest to determine the amount of fibroblasts, tumor cells, and immune cells via cell surface marker staining or GFP expression, as indicated.

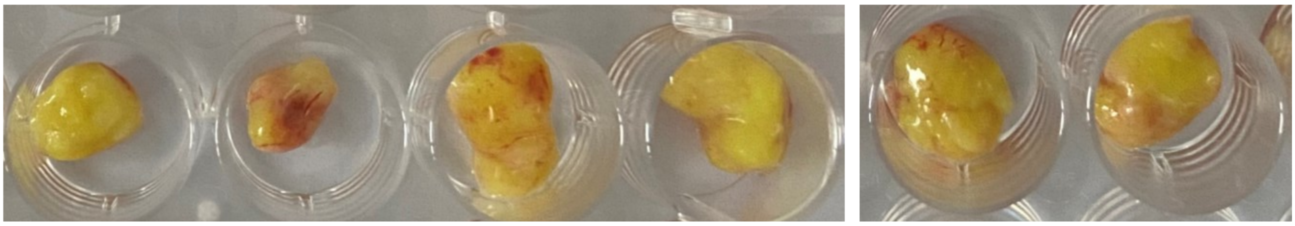

**FAP-Ad5-TZB** **PBS** **Herceptin 1x**

**Supplementary Figure 5: Delivery of TZB via FAP-Ad5 is more effective in reducing tumor growth than direct injection of the recombinant protein.** NCI-N87 tumor cells and NIH3T3mFAP cells were co-injected subcutaneously into the flank of SCID/beige mice. At a tumor volume of 50 $mm^3$, mice were treated intratumorally with $9 \times 10^8$ PFU FAP-Ad5-TZB (n = 5), or 200 µg Herceptin (n = 3), or PBS (n = 5). Tumors were harvested 19 days post-injection and analyzed further. Two representative samples per group are depicted here.
